# Resource-efficient nitrogen removal from source separated urine with partial nitritation/anammox in a membrane aerated biofilm reactor

**DOI:** 10.1101/2023.12.21.572732

**Authors:** Marijn J. Timmer, Jolien De Paepe, Tim Van Winckel, Marc Spiller, Irina Spacova, Kim De Paepe, Isabel Pintelon, Sarah Lebeer, Winnok H. De Vos, Christophe Lasseur, Ramon Ganigué, Kai M. Udert, Siegfried E. Vlaeminck

## Abstract

Source separation and decentralized urine treatment can cut costs in centralized wastewater treatment by diverting 80% of the nitrogen load in sewage. One promising approach for nitrogen removal in this context is partial nitritation/anammox (PN/A), reducing the aeration demand by 67% and organics dosage by 100% compared to nitrification/denitrification. Whilst previous studies with suspended biomass have encountered stability issues during PN/A treatment of urine, a PN/A biofilm was hypothesized to be more resilient. Its use for urine treatment was pioneered here for maximum rates and efficiencies in the energy efficient membrane-aerated biofilm reactor (MABR). Nitrogen removal rates of 1.0 g N L^-1^ d^-1^ and removal efficiencies of 80-95% were achieved during a 335-day stable operation at 28°C on stabilized (pH>11), diluted urine (10%). A balance between N_2_ and NO_3_ ^-^ formation was observed whilst optimizing the supply of O_2_ and was rate limiting for the conversion towards N_2_. Short-term operation on less- and undiluted urine yielded N removal rates of 0.6-0.8 g N L^-1^ d^-1^ and removal efficiencies of 93% on 66% urine and 85% on undiluted urine. Metataxonomic analysis and fluorescence *in-situ* hybridization confirmed the presence of biofilms consisting of nitrifiers (*Nitrosomonas, Nitrospira*) at the membrane side and anammox bacteria (“*Candidatus* Brocadia”) at the anoxic bulk side. The findings suggest that a biofilm approach to PN/A treatment of urine overcomes stability issues, and a PN/A-MABR has significant potential for resource efficient decentralized treatment. In human long-duration deep-space missions, this gravity-independent technology could produce N_2_ to compensate artificial atmosphere losses whilst facilitating water recovery from urine.

**[GRAPHICAL ABSTRACT, COLOR]:** 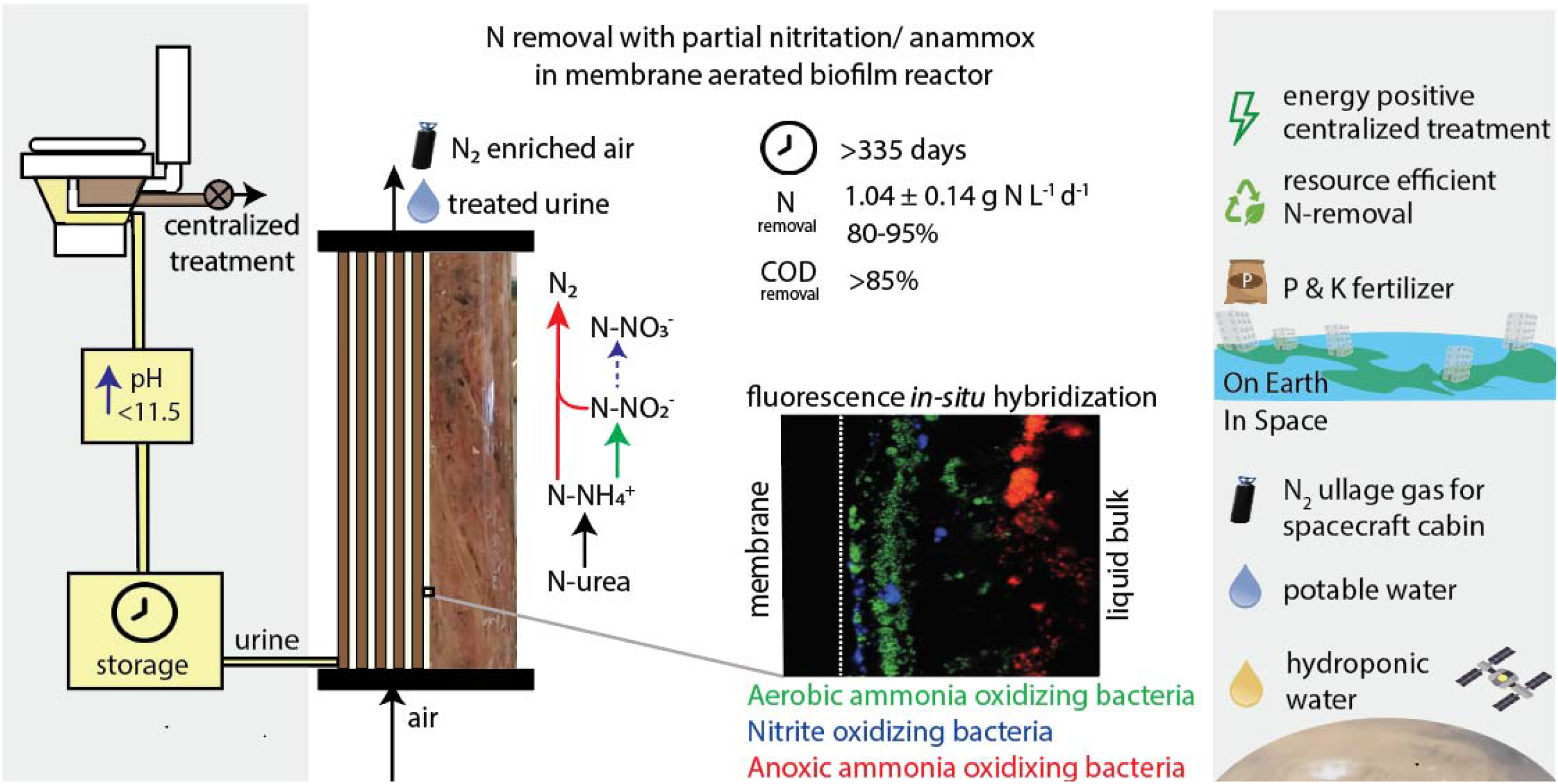

## 1. Introduction

Reactive (organic and ammoniacal) nitrogen is one of the main constituents in domestic wastewater and is typically removed biologically through nitrification/denitrification (N/DN) in centralized wastewater treatment plants (WWTP). Nevertheless, N/DN represents a large fraction of the operational costs of WWTP because of the energy-intensive aeration (45 MJ kg^-1^ N) (Maurer et al., 2003). Roughly 75-80% of the nitrogen in domestic wastewater originates from urine, while urine accounts for less than 1% of the total volume (McConville et al., 2017). Therefore, the separate collection of urine and faeces at the toilet (source separation) has gained considerable attention as an energy-efficient alternative to the existing end-of-pipe wastewater management (Larsen et al., 2021). A separation efficiency of 60-75% of the urinary nitrogen would eliminate the need for N/DN, as all nitrogen would be used for assimilation (Wilsenach and van Loosdrecht, 2003). This would reduce operational complexity and even make wastewater treatment energy neutral (Siatou et al., 2020). In addition, the source-separated urine can be processed with a fit-for-purpose technology, e.g. nitrification to produce a redox buffer in the sewer (Jiang et al., 2011), (half) nitrification to produce fertilizer for plants or microalgae (Coppens et al., 2016; Udert and Wächter, 2012), or nitrogen removal with resource-efficient partial nitritation/anammox (PN/A). PN/A requires 60% less oxygen compared to N/DN, hence requires significantly less aeration energy (19 MJ kg N^-1^, (Maurer et al., 2003)). Moreover, it is an autotrophic process, which eliminates the need for additional chemical oxygen demand (COD) dosage to denitrify urine. PN/A on urine involves a three-step process. First, urine is hydrolyzed to ammonia by heterotrophs. To prevent ammonia volatilization and odour emission, urine prestabilization can be performed at cost of energy (Paepe et al., 2020) or chemicals (Randall et al., 2016).

Ammonia is then partially oxidized to nitrite by aerobic ammonia oxidizing bacteria (AerAOB). Subsequently, ammonia and nitrite are converted to nitrogen gas by anoxic ammonia oxidizing bacteria (AnAOB). One of the key operational challenges of the PN/A process is to suppress the potential oxidation of nitrite to nitrate by nitrite oxidizing bacteria (NOB), as this increases the oxygen consumption and decreases the N removal efficiency. Different strategies have been reported to suppress NOB activity such as leveraging free ammonia (FA), free nitrous acid and intermittent aeration (Van Tendeloo et al., 2021), but have never fully succeeded to eliminate it. The use of a combination of free ammonia residual with precise aeration control could yield better NOB suppression.

Besides the need for NOB suppression, the application of PN/A for urine treatment presents further challenges due to its intricate matrix, particularly with limited dilution in source-separated systems (2-4 g N L^-1^). Fresh or stabilized urine has an average conductivity of 20-30 mS cm^-1^ which poses a challenge for the proliferation of AnAOB. Furthermore, the high mono/divalent cation ratio in urine can lead to partial inhibition and affect floc formation in flocculent systems (Dapena-Mora et al., 2010), with subsequent washout. To date, PN/A has only been shown on diluted (10-20%) urine with lab-scale sequencing batch reactors as suspended growth systems and achieved limited success due to AnAOB inhibition and washout (Bürgmann et al., 2011). To the best of the authors’ knowledge, PN/A treatment on undiluted urine has never been successfully demonstrated, potentially due to the inhibitory effects of salinity, organic compounds, and variations in COD/N ratio (Schielke-Jenni, 2015).

A potential solution to the outlined challenges is the use of a membrane-aerated biofilm reactor (MABR), which offers an energy-efficient aeration alternative that can save up to 70% in aeration costs compared to conventional bubble aeration (Syron et al., 2015). Simultaneously, a high membrane packing density (300-500 m^2^ m^-3^) (Castrillo et al., 2019; Syron and Heffernan, 2017) allows for small system footprint. MABRs enable precise oxygen supply and high N and COD removal rates in simultaneous oxic/anoxic systems (Downing and Nerenberg, 2008) as they uncouple substrate and oxygen diffusion with counter-diffusion. In contrast, biofilm systems with bubble aeration that rely on co-diffusion do not provide the same level of control over dissolved oxygen (DO), limiting the ability to manage NOB growth, promote AnAOB activity in the biofilm, and effectively control the ratio of oxic and anoxic zones in reactors that experience fluctuating COD/N ratios, such as urine treatment systems.Besides this, the biofilm instead of suspended flocs could overcome the stability issues one urine as they are known for their ability to produce extracellular polysaccharide substances (EPS) that provide a protective barrier against fluctuations in (potentially) toxic compounds, salinity, and N levels, etc… Biofilm systems have demonstrated the ability to overcome washout-effects and operate successfully in high saline conditions, even up to seawater, 50-60 mS cm^-1^ (Kartal et al., 2006; Windey et al., 2005). However, the improved resilience on a urine matrix has never been investigated. Consequently, PN/A in MABR was widely reported successfully in lab scale on different streams (Dong et al., 2009; Gong et al., 2007; Sun et al., 2009) even on full scale (Bunse et al., 2020). Whilst nitrification is already tested in a lab-scale MABR on urine (Paepe et al., 2020), the use of PN/A was only reported once and obtained low rates, uncompetitive to N/DN (up to 0.4 g N L^-1^ d^-1^) (Zhan et al., 2022) on diluted urine, whilst the technology has never been established on undiluted urine.

We hypothesized that the ease of biomass retention and the resilience of a biofilm system against process fluctuations, in combination with interval membrane aeration could yield high rate, stable autotrophic N removal from urine. To test this hypothesis, this study investigated the feasibility of resource-efficient nitrogen removal from urine via PN/A in an MABR. Our primary aim was to reach maximum N and COD removal rates and efficiencies trough optimized intermittent aeration and feeding on 10 times diluted urine via the energy efficient PN/A pathway. Our secondary aim was to assess the capacity of the MABR to overcome inhibitory effects of undiluted urine observed in other reactor types. Thirdly, the study aimed to describe the dynamics and spatial distribution of AerAOB, NOB and AnAOB in a counter-diffusional biofilm treating urine.

## 2. Materials and Methods

### 2.1. MABR set-up

The MABR was composed of 3 or 4 hollow fibre bundles (Figure 1), each consisting of 180 dense (non-porous) silicone rubber hollow fibres with a length of 25 cm, an inner diameter of 0.3 mm and an outer diameter of 0.55 mm (Nagasep M100, Nagayanagi Co., Saitama, Japan). Each bundle had a total membrane surface area of 0.06 m^2^ and liquid volume of 0.09 L (area to volume ratio of 667 m^2^ m^-3^). The total system volume varied between 0.57 L (3 bundles) and 0.72 L (4 bundles) throughout the study (Table 1). Liquid was recirculated over the shell side of the hollow fibres with a peristaltic pump (323 D/E, Watson Marlow, Marlow, UK) (150 mL min^-1^ per bundle). Air was humidified to saturation using washing bottles and transported through the lumen of the fibres using gas pumps (APS 100, Tetra, Melle, Germany) with a flow rate of 0.5 L min^-1^ co-current to the liquid. The DO was monitored online (interval 1 sec) using a DO-sensor (InPro 6050, Mettler Toledo, Columbus, USA) and controller (M300, Mettler Toledo, USA). The pH was controlled between 7 and 7.5 by acid addition (0.25M HCl) (Consort R3610, Turnhout, Belgium). The temperature was controlled at 27.5 ^⍰^C with a thermostat (ITC308, Inkbird, Shenzhen, China) coupled to a heating cable.

**Table 1.**
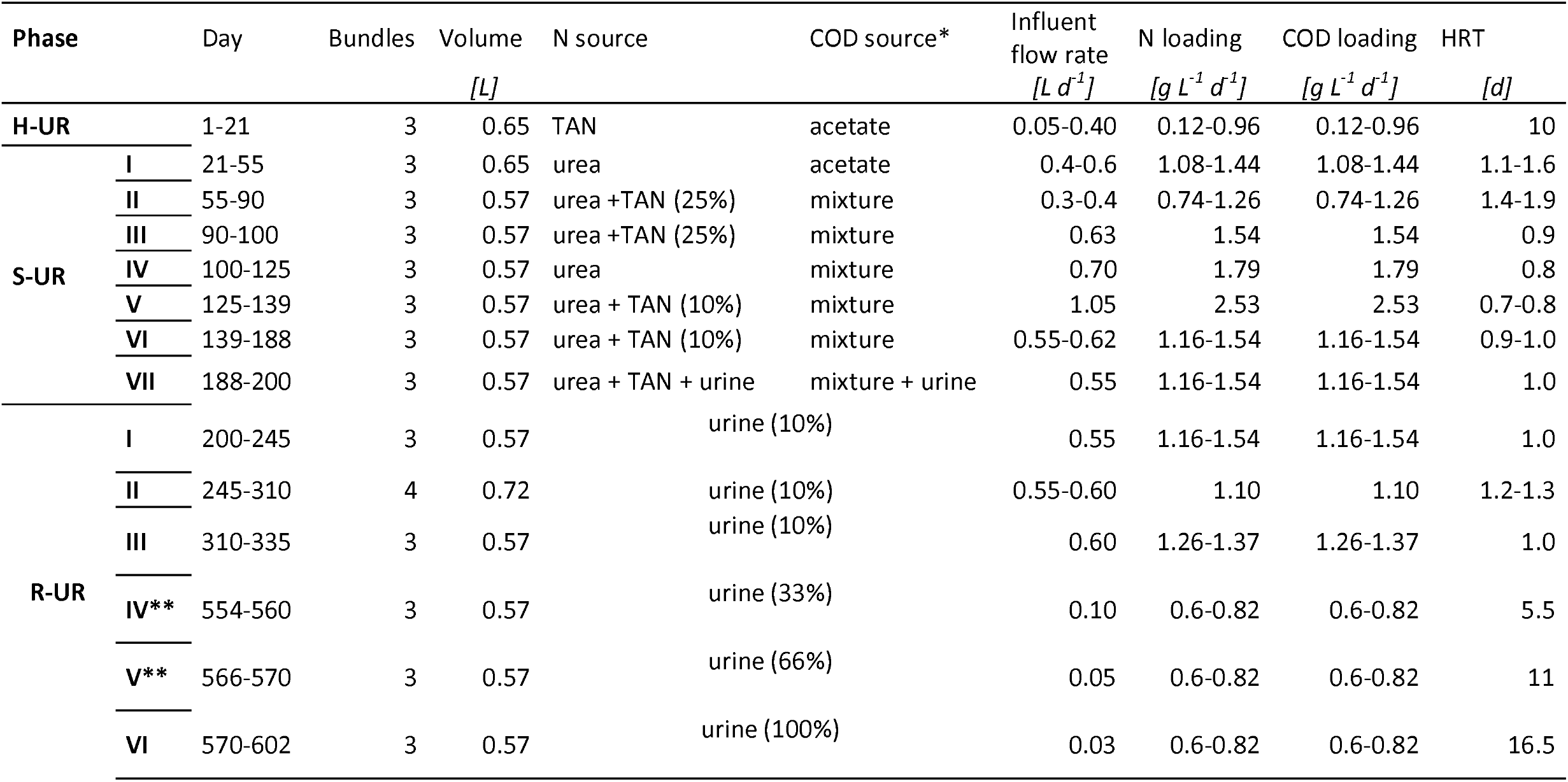
Phases of reactor operation. HUR: (synthetic) hydrolysed urine, S-UR: synthetic urine, R-UR: real urine. * Mixture: mixture of creatinine (0.462 g L^-1^ ), hippuric acid (0.140 g L^-1^) and citric acid (0.168 g L^-1^). ** No steady state was obtained.

**Figure 1.**
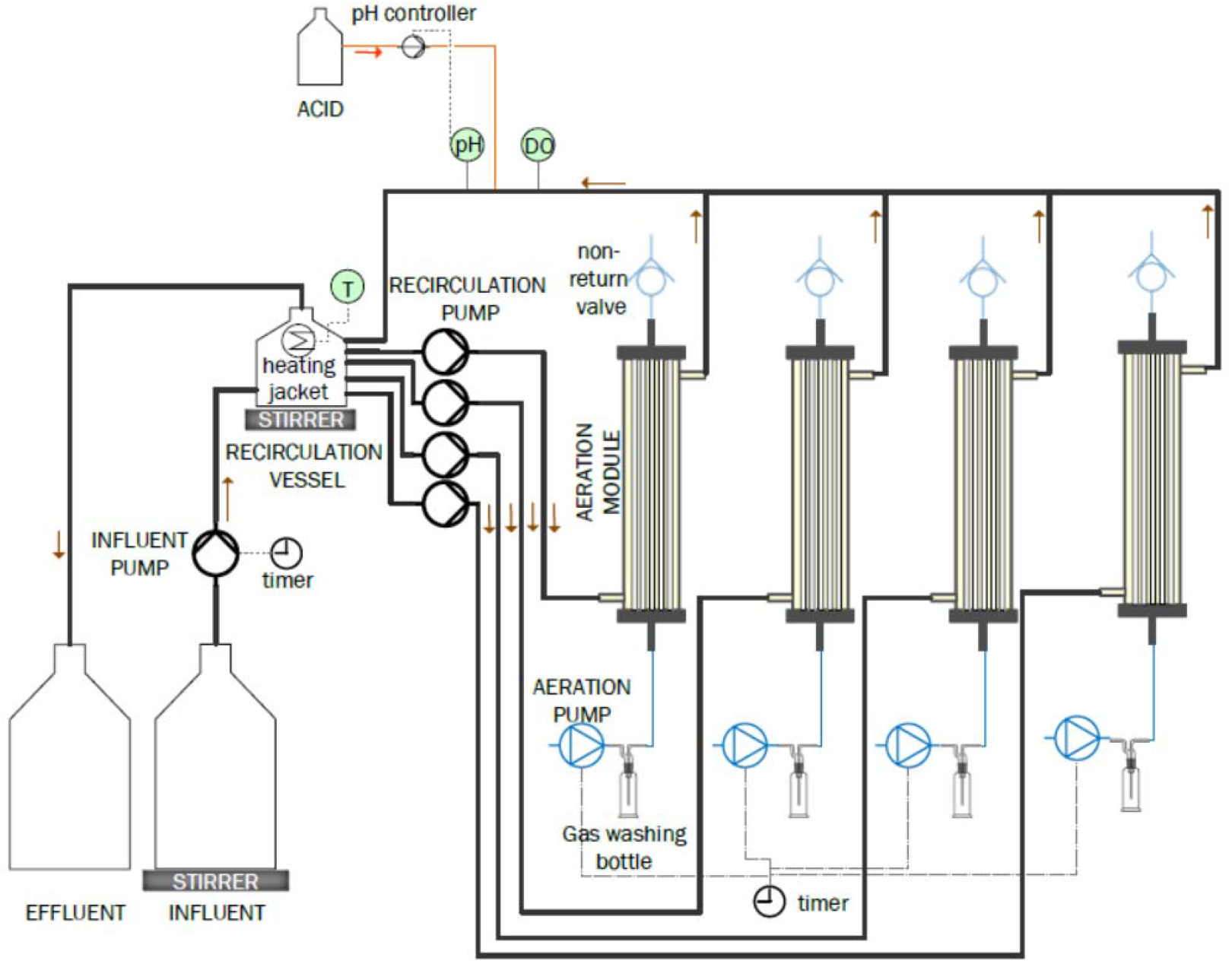
Schematic overview of the PN/A-MABR. The here displayed configuration is composed of 4 bundles, although throughout the study, the reactor was operated with 3 or 4 bundles.

The influent pump (323 D/E, Watson Marlow) was connected to a timer (SYN 151h, Theben, Haigerloch, Germany) and interval dosing with a dosage velocity of 0.6-50 mL h^-1^ in different operation phases (Table 1). The influent was continuously stirred to prevent settling of precipitates that formed during alkanization, mostly composed of magnesium- and calcium and phosphate. Effluent left the system by a pressure-based overflow mechanism.

### 2.2. MABR operation

The MABR was inoculated with 19 g VSS L^-1^ PN/A sludge from a lab-scale rotating biological contactor operated on synthetic wastewater (recipes in SI Table S2) (Van Tendeloo et al., 2021). To avoid biomass starvation from limited urea hydrolysis, the MABR was first operated on diluted synthetic hydrolysed urine (0.67 g N L^-1^). After 21 days, the reactor was transferred to synthetic urine, which contained urea instead of TAN as N-source (phase S-UR I, Table 1), which was reversed partially in subsequent operational phases to resolve ureolytic bottlenecks (SUR II-III). In phase S-UR II-VI, a COD-mixture composed of the three major COD compounds in urine (Putnam, 1971) was used instead of sodium acetate. Finally, in phase S-UR VII a mixture of half synthetic and half real urine was used as transition to the real urine. All media were alkalinized to pH >11 with 2M NaOH to mimic the pH of the stabilized urine. A concentration of 670 mg N & COD L^-1^ was used in all recipes, corresponding to a ten times diluted urine matrix with a COD/N-ratio of 1. A bioaugmentation was done after 120 days with a volatile suspended solid (VSS) concentration of 70 mg L^-1^ from a high-rate activated sludge unit treating sewage and 100 mg L^-1^ VSS from a PN/A unit treating sludge reject water (WWTP Nieuwveer, Breda, the Netherlands) in response to the observed ureolytic limitation.

After 200 days of operation, the reactor was switched to real urine (R-UR), collected from male donors at our facility (urine collection was approved by the Ethical Committee UZA-UA, registration number B3002021000034). Urine donations were pooled into batches of 4 L. The urine was immediately stabilized to pH<11 with 2M NaOH. Urine was stored up to 8 weeks at room temperature and diluted with tap water in smaller batches prior to feeding to the reactor. The total nitrogen (TN) of the urine was measured to determine the dilution needed to obtain a concentration of 670 mg N L^-1^. The total ammonia nitrogen (TAN)-concentration and the pH were checked before dilution to ensure stability of the stored urine.

From day 554, the urine fraction in the medium was stepwise increased from a ∼10% (0.67 g N L^-1^) to a ∼100% (5.5 g N L^-1^) urine solution, the maximum we could obtain from our collection at the facility (probably due to the lack of morning urine). Two intermediate steps of 2 g N L^-1^ and 4 g N L^-1^ were used, at a constant loading rate of 0.6-0.8 g N L^-1^ d^-1^, but due to the increased hydraulic residence time (HRT), no steady state was observed at the intermediate phases (R-UR IV-V). Feeding and aeration control is described in (SI Table S1). Influent and effluent samples were taken every 1-3 days and were filtered with a 0.2 μm filter before being stored at 4 ^⍰^C prior to analysis.

Sludge was harvested from the reactor to prevent reactor clogging. This was done based on visual observation and performed on days 127, 202 and 335 by scouring the bundles. On day 245, a fourth membrane bundle was added to the system and cross-inoculated from the other membrane bundles in anticipation of the sacrifice of another bundle on day 310 for community analysis and FISH. During scouring events, 0.7-1.7 g VSS was removed, representing 5-12% of total 12.52 g VSS (in 3 bundles, based on estimation after bundle sacrifice).

### 2.3. Analytical methods

Anion concentrations were analyzed on an Ion Chromatograph (Metrohm 930 with Metrosep A supp 5-150/4.0 column, Herisau, Switzerland). TAN concentrations were measured using a San++ Automated Wet Chemistry Analyzer (Skalar, Breda, The Netherlands). Test kits TN 220 and COD 1500 (Macherey-Nagel, Pennsylvania, USA) were used to measure the TN and COD concentration. Electrical conductivity (EC) was measured with a conductivity meter (HI2030-02a, Hanna Instruments, Woonsocket, USA). pH was measured with a portable pH meter (HI2020-02, Hanna Instruments). Indirect measurements for nitrogen gas produced and organic nitrogen were derived considering a mass balance over the TN in- and effluent and subtraction of TAN, NO_2_^-^, and NO_3_^-^.

### 2.5 16S rDNA amplicon sequencing of microbial communities

For the taxonomic microbial community analysis, sludge samples were centrifuged, and the resulting pellets were stored at -20^⍰^C before DNA extraction. DNA was extracted using the PowerSoil (Qiagen, Hilden, Germany) extraction kit according to the protocol from the manufacturer. The V3-V4 region of the 16S rRNA gene was amplified with 341F-806R universal primers derived from (Klindworth et al., 2013) and (Caporaso et al., 2011), and sequenced on a Novaseq platform generating 250 bp paired end reads (Illumina, San Diego, California, US) by Novogene (Beijing, China). The pipeline used for processing the resulting Miseq data is described in Supplementary Information, D.

### 2.4. Fluorescence *in situ* hybridization analysis of bacterial stratification in biofilms

To study the stratification of AerAOB, NOB and AnAOB, fluorescence *in situ* hybridization (FISH) was performed on cryosections of the biofilm. Representative membrane sections of 2 cm were harvested from the middle of the sacrificed bundle (to exclude heterogeneity at the inflow or outflow part). Twenty samples were fixed according to (Nielsen et al., 2009) with (4%) paraformaldehyde buffer and an incubation time of 6 h. Subsequently, samples were stored in 1:1 phosphate-buffered saline (PBS): ethanol at -20^⍰^C. After washing in PBS and overnight incubation in PELCO^®^ Cryo-Embedding Compound (Ted Pella Inc., Redding, USA) at room temperature(RT), samples were frozen at -20^⍰^C and cryosections with 30 μm thickness were prepared Leica CM1950 cryomicrotome (Leica BioSystems, Diegem, Belgium). Slides were then thawed on Superfrost glass slides (Epredia, Porthsmouth,USA), stepwise dehydrated with 50, 70 and 96% ethanol, and stored at RT. *In situ* hybridization was performed using fluorescent probes from Eurogentec (SI Table S3) in two hybridization rounds at 40% and 15% formamide hybridization buffers according to (Vlaeminck et al., 2009) and a 4⍰,6-diamidino-2-phenylindole(DAPI) staining was performed according to (Nielsen et al., 2009). Samples were immersed in antifadent Vectashield (Vector laboratories, Burlingame, USA) to prevent fluorophore degradation. Images were obtained using a Leica DMi8 inverted fluorescence microscope (Leica Microsystems, Diegem, Belgium) equipped with a 405-nm diode laser (to detect DAPI) and a white light laser (WLL) used at 488, 555 and 635 nm to visualize respectively FITC, Cy3 and Cy5. Images are single confocal planes. Images were acquired at magnifications of 100,630 and 950x with a resolution of [1024 x 1024] pixels.

## 3. Results

### 3.1. Maximum nitrogen and COD removal rate on diluted urine

After a start-up phase of 21 days on synthetic hydrolysed urine with only TAN as N-source (Figure 2A), the system was transferred to synthetic urine (phase S-UR I), where several limitations emerged that prevented optimal PN/A performance. An increasing nitrate (>0.1 g N L^-1^, Figure 2B) residual was observed in the first 50 days (S-UR I and II). The nitrate formed originated from NOB activity as nitrate originating from anammox metabolism (11% of N converted) can be denitrified using the COD in urine. To suppress NOB, a strict time based instead of DO-aeration regime was implemented of initially only 10 min h^-1^ (Figure 3A) which eliminated the nitrate residual (S-UR II). Ureolytic bottlenecks were indicated by an organic nitrogen residual in the effluent after the change to urea as main nitrogen source (S-UR II-IV), indicating incomplete. This was addressed by first changing back to 25% TAN (S-UR-II,III) instead of only urea (Figure 2A), then by introducing a mixture of creatinine, hippuric and citric acid as COD instead of acetate increase diversity in the community, and finally by bioaugmentation (S-UR IV), of which only the last strategy effectively enhanced organic nitrogen removal. Consequently, a TAN-residual up to 0.3 g N L^-1^ then formed (S-UR V-VII), associated with FA levels of 7 mg N L^-1^ . As mitigation for the TAN residual the aeration was stepwise increased to 25 min h^-1^ (S-UR IV-VI) to enhance nitritation. Simultaneously the loading rate was increased to 2.6 g N L^-1^ d^-1^ . This increase in loading rate did not lead to an increase of the TAN-residual in the effluent, whilst a maximum nitrogen removal rate (NRR) of 1.6 g N L^-1^ d^-1^ (2.5 g N m^-2^ d^-1^, SI Figure S3) between day 130 and 140 could be reached at a loading rate of 2.6 g N^-1^ L^-1^ d^-1^ (Table 2). The FA effectively supressed NOB activity, reflected by the NO_3_ levels of 0 mg N L^-1^ . Nevertheless, the high TAN residual resulted in a nitrogen removal efficiency (NRE) of only 60-70%. The maximum NRR of 1.6 g N L^-1^ d^-1^ is higher than the maximum anammox activity of 1.45 g N L^-1^ d^-1^ found with anoxic activity tests (SI Table S4), which is the result of denitrification occurring in the reactor, whilst we did not add COD during the activity tests. However, the proximity to the maximum anammox activity indicates that the performance was limited at this point by both aeration and anoxic activity.

**Table 2.** Summary of performance parameters at maximum nitrogen removal rate and efficiency achieved for the PN/A-MABR treatment of 10 times diluted urine.

| Parameter | Unit | Maximum nitrogen removal rate | Maximum nitrogen removal efficiency |
| --- | --- | --- | --- |
| Period | [d] | 130-140 | 250-260 |
| Nitrogen Loading Rate | [g N L<sup>-1</sup> d<sup>-1</sup>] | 2.4 | 1.1 |
| NRR (Volumetric) | [g N L<sup>-1</sup> d<sup>-1</sup>] | 1.6 | 1.0 |
| NRR (Area) | [g N m<sup>-2</sup> d<sup>-1</sup>] | 2.5 | 1.5 |
| NRE | [%] | 60-70 | 80-95 |
| NRR | [g TKN-N L<sup>-1</sup> d<sup>-1</sup>] | 1.6 | 1.1 |
| COD-RE | [%] | 85 | 85 |

**Figure 2.**
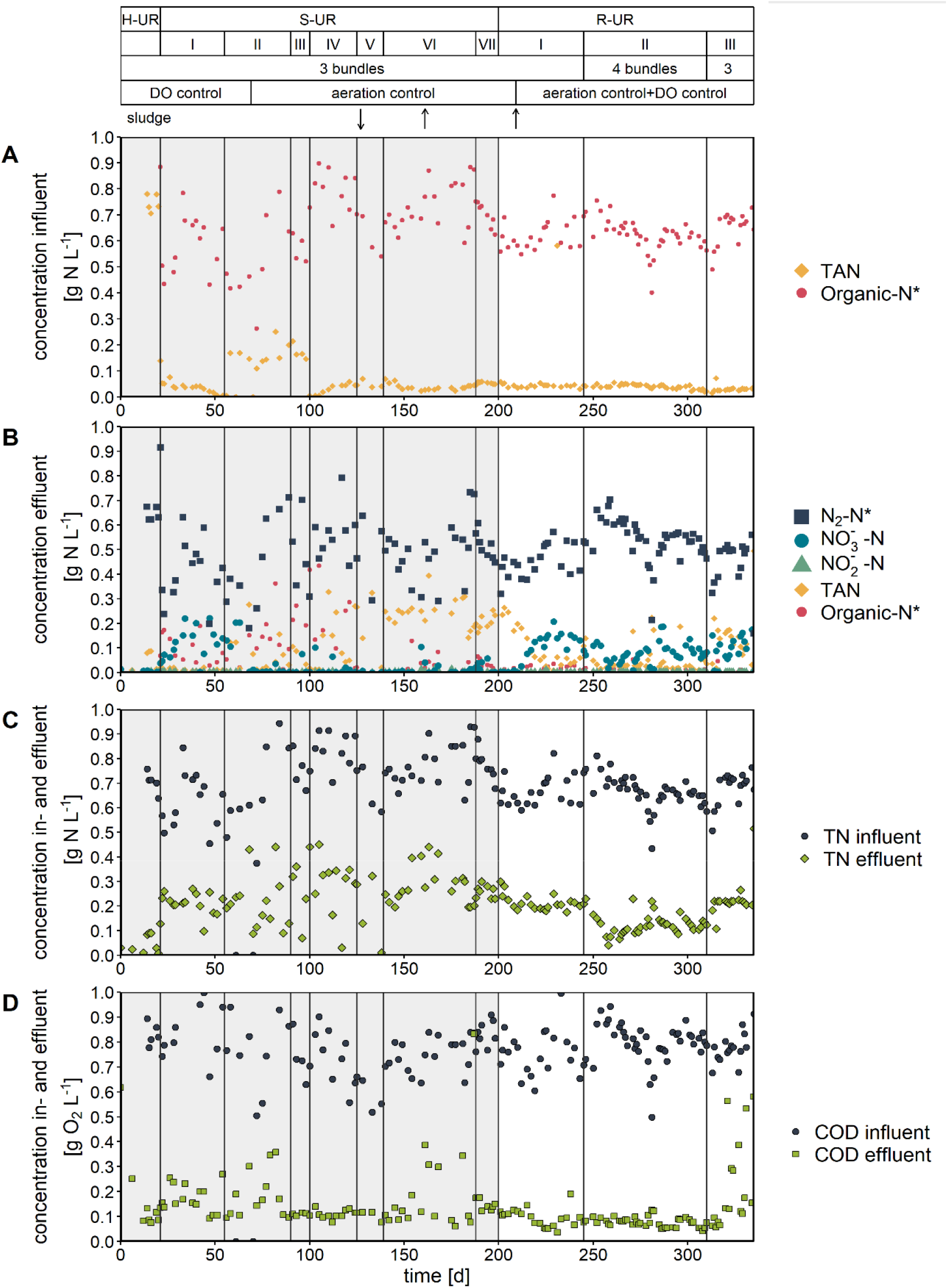
Concentrations N and COD in MABR in- and effluent during reactor operation on synthetic (grey) and real (white) diluted urine. Nitrogen speciation in the influent (A) and effluent (B), total nitrogen in influent and effluent (C), and COD concentration in influent and effluent (D). In the top legend, sludge arrows down resp. up represent a bioaugmentation resp. sludge harvesting event *calculated from indirect measurements. HUR: (synthetic) hydrolysed urine, S-UR: synthetic urine, R-UR: real urine.

**Figure 3.**
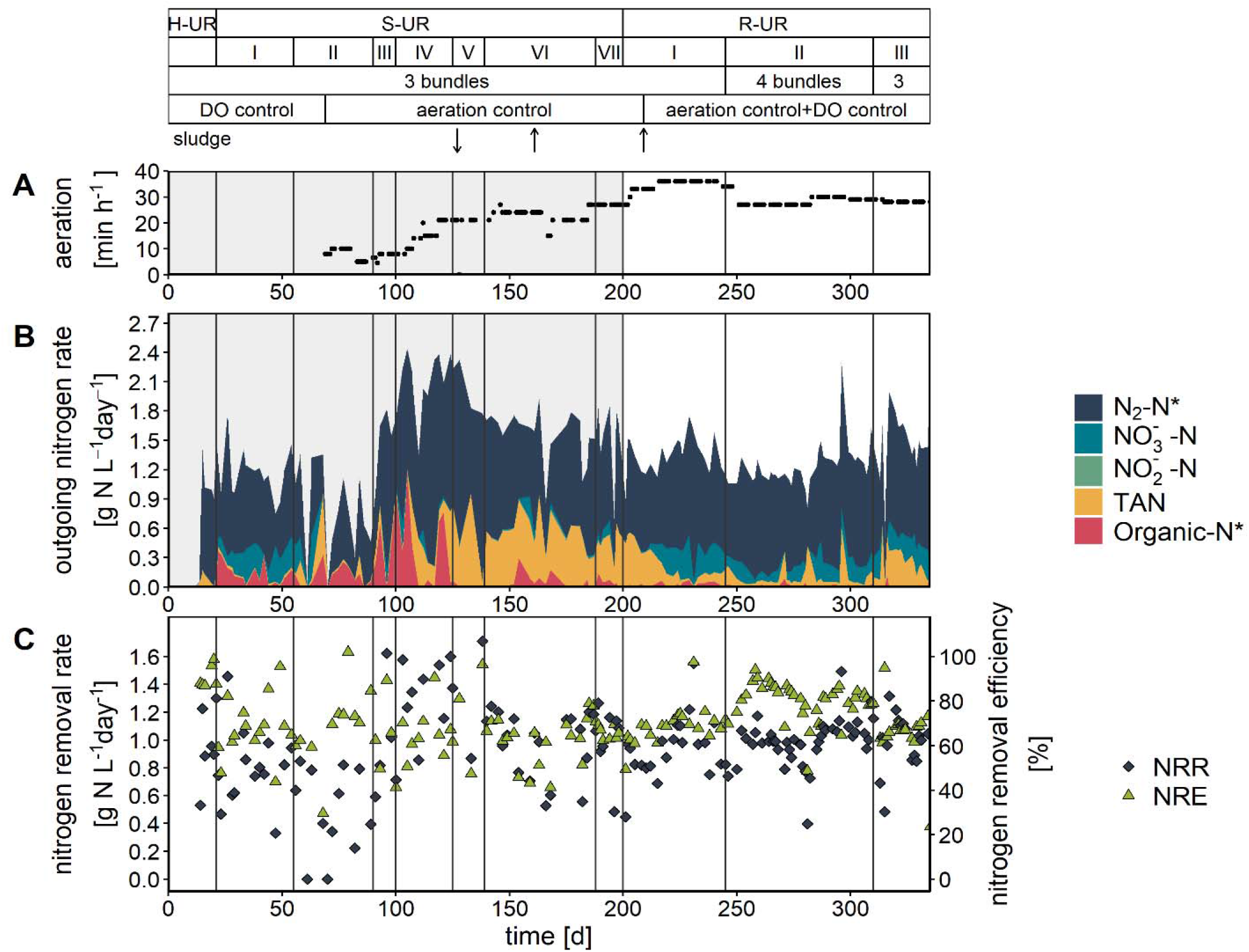
Nitrogen fate, removal rate and efficiency during reactor operation on synthetic (grey) and real (white) diluted urine. A) Minutes per hour of aerated time B) Speciation of outgoing nitrogen per day C) Nitrogen removal rate and nitrogen removal efficiency. In the top legend, sludge arrows down resp. up represent a bioaugmentation resp. sludge harvesting event *calculated from indirect measurements. HUR: (synthetic) hydrolysed urine, S-UR: synthetic urine, R-UR: real urine

### 3.2. Maximum removal efficiency on real diluted urine

Once the startup was established, several bottlenecks were overcome and a high PN/A activity was achieved, the focus shifted to proofing the process on real urine (S-UR VII, day 190-200) and optimizing the NRE. To achieve this, the TAN-residual (up to 0.3 g N L^-1^) had to be reduced. The loading rate was lowered to 1.0 g N L^-1^ (S-UR VI) and the aeration time was further increased to 30 min h^-1^, but both changes did not yield in significant improvement of NRE (Figure 3C) as the TAN-residual remained 0.2-0.25 g N L^-1^ (S-UR VI-VII). At day 189, the switch from synthetic to real urine (S-UR VII) was made, which did not affect concentrations of N and COD in the effluent, remaining below 0.25 g N L^-1^ (mostly as TAN) and 0.15 g COD L^-1^ respectively (Figure 2, B&D).

In contrast to previous aeration time increases, when aeration time was increased up to 35 min h^-1^, significantly lower TAN concentrations (>0.05 g N L^-1^ ) were observed (R-UR I). Yet, the TAN-residual was merely replaced by NO_3_ ^-^(0.15 g N L^-1^ ), caused by NOB activation in the absence of a TAN residual. As a result, NRE increased only from 60 to 70%. To prevent NOB activity following from a residual DO, during the aerated time, a DO-shutoff was added to the timer based oxic/anoxic control (R-UR I), which led to stabilization of the amount of NO_3_^-^ formed (0.15 g NL ), but not to suppression. Upon the addition of a fourth, clean membrane bundle, the NRE increased to its maximum at 90% between day 250-260 at a NRR of 1.0 g N L^-1^ d^-1^ (1.5 g N m^-2^ d^-1^), outlining the trade-off between optimum NRR or NRE (Table 2). The initial absence of a biofilm in the clean bundle might have led to a more precise control of diffusion in the liquid bulk, where the DO was measured, leading to the optimum NRE. After day 260, the NRE started to decrease as a NO_3_ ^-^residual (up to 0.1 g N L^-1^ ) formed. In response to the NO_3_^-^residual, the aerated time was lowered again at day 309, but this had no effect on the NO_3_^-^ formation rate. Hence, even with a timer-based oxic anoxic control strategy supplemented with a DO-control, it remained challenging to supply the right amount of oxygen the prevent TAN and NO_3_ accumulation. The data suggest that thicker biofilms had worse controllability, the DO-shutoff was less effective due to diffusion limitation, and allowed for more NOB to grow. As opposed to the time (min h^-1^ ) the system was aerated, the interval and spread of the aeration per hour were not found to impact the performance (SI Table S1).

NRR and NRE did not seem to impact the removal of COD, which remained consistently stable, at approximately 85% for both synthetic and real urine (Figure 2D).

### 3.3. Nitrogen removal rate and efficiency on undiluted urine

In the last operation phases of the reactor, PN/A in MABR on less diluted and undiluted urine was investigated( R-UR IV-VI, day 554-595) with TN-concentrations from 0.67, 2, 4 and 5.5 g N L^-1^ (Figure 4E). A loading rate lower than on operation on diluted urine of 0.7 g N L^-1^ d^-1^ was set to avoid a crash due to overloading/adaptation effects. Whilst NRRs of 0.6 g N L^-1^ were reached for all dilutions, the NRE varied for the operation on different urine concentrations (Figure 4C). An optimum in NRE of 92% was achieved at a dilution of 66% (4 g N L^-1^ ), but this condition was only run for a short operation span (5 d). Although steady state performance was therefore not achieved, the transition from 66% to 100% undiluted urine hinted even at a further increase in NRE (95%, SI Figure S4), indicating that no inhibitory effects for PN/A were observed at effluent EC similar to 66% influent urine(<20 mS cm^-1^ ). However, when the EC reached 20 mS cm^-1^, NRE lowered to 85% when steady state was approached (HRT 16.5 d). At this point, a TAN and NO_3_^-^ residual of 0.2 and 0.5 g N L^-1^ was formed, hinting at partial inhibition of both AerAOB and AnAOB, most likely caused by the high salinity as both aeration and loading were not changed throughout R-UR VI-VII. AnAOB activity tests conducted at different strengths of urine matrix confirmed the partial inhibition effect on AnAOB (SI Figure S5), outlining a drop in the maximum activity of 33% with 100% urine compared to 10% urine. Given that only 2 HRTs were operated for the 100% urine, it remains to be observed if these rates and efficiencies on 100% urine can be maintained for long-term steady state operation.

**Figure 4.**
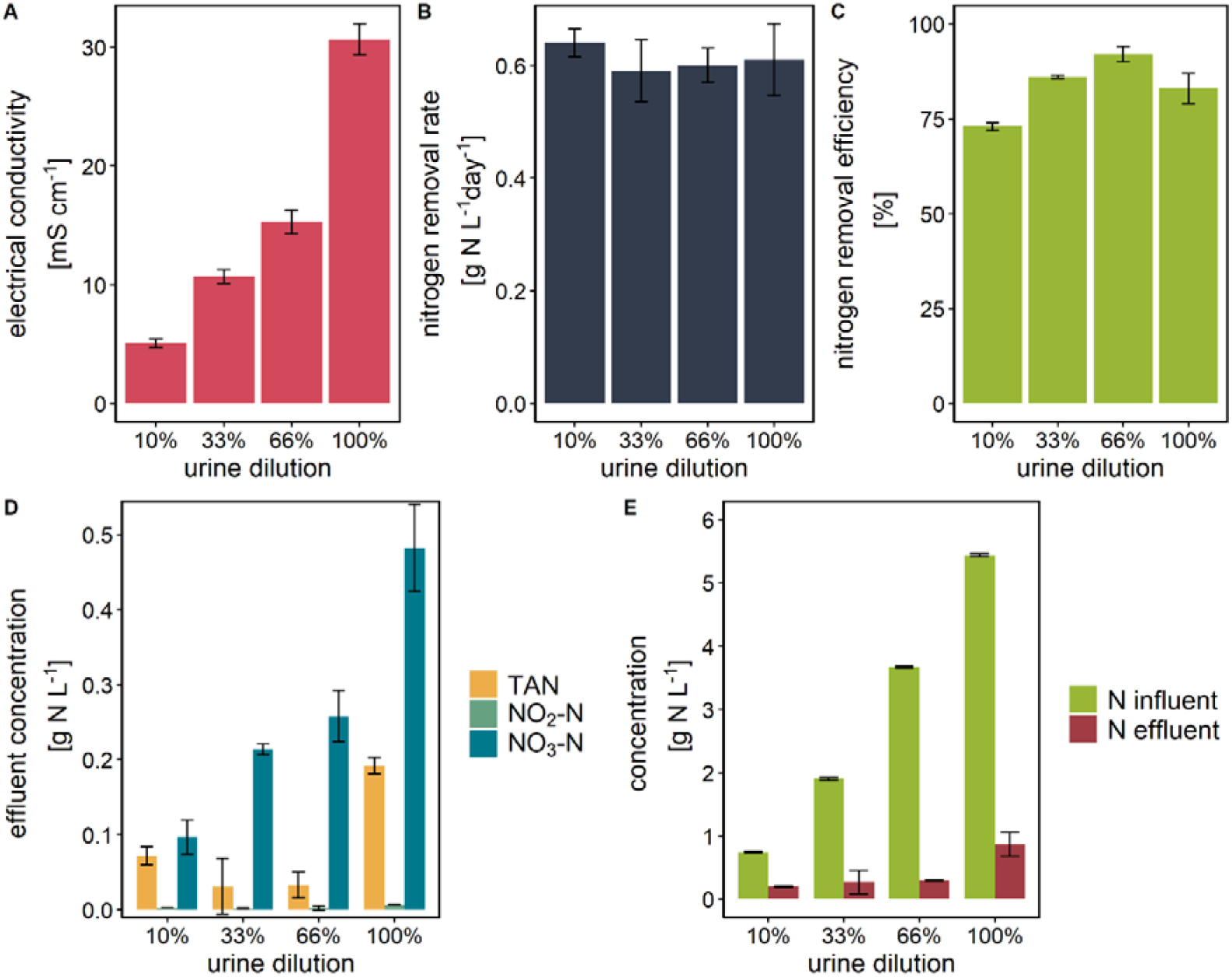
Reactor performance on 10%, 33%, 66% and 100% real urine. A) Electrical conductivity in the effluent B) Nitrogen removal rate. C) Nitrogen removal efficiency D) Concentration of TAN, NO_2_^-^ -N and NO_3_^-^ -N in the effluent E) Concentration total nitrogen in in- and effluent. Although the concentrations were increased, the loading rate was kept the same.

### 3.4. Taxonomic microbial 16S rDNA community analysis

The microbial community composition was monitored over time to investigate a potential link between performance of PN/A on urine and microbial community. Related to nitrogen removal, *Candidatus* Brocadia from the *Brocadiacea* family was the only AnAOB making up 1-5% of the total identified community during operation on synthetic and real urine, as opposed to the inoculum where it made up 14% of the community (Figure 5B,D). *Nitrosomonas* (family *Nitrosomonadaceae*) followed a similar trend in abundance and was the only identified AerAOB. As for NOB, *Nitrospira* (family *Nitrospiraceae*) and *Nitrobacter* (family *Nitrobacteraceae*) were identified, *Nitrospira* being dominant.

**Figure 5.**
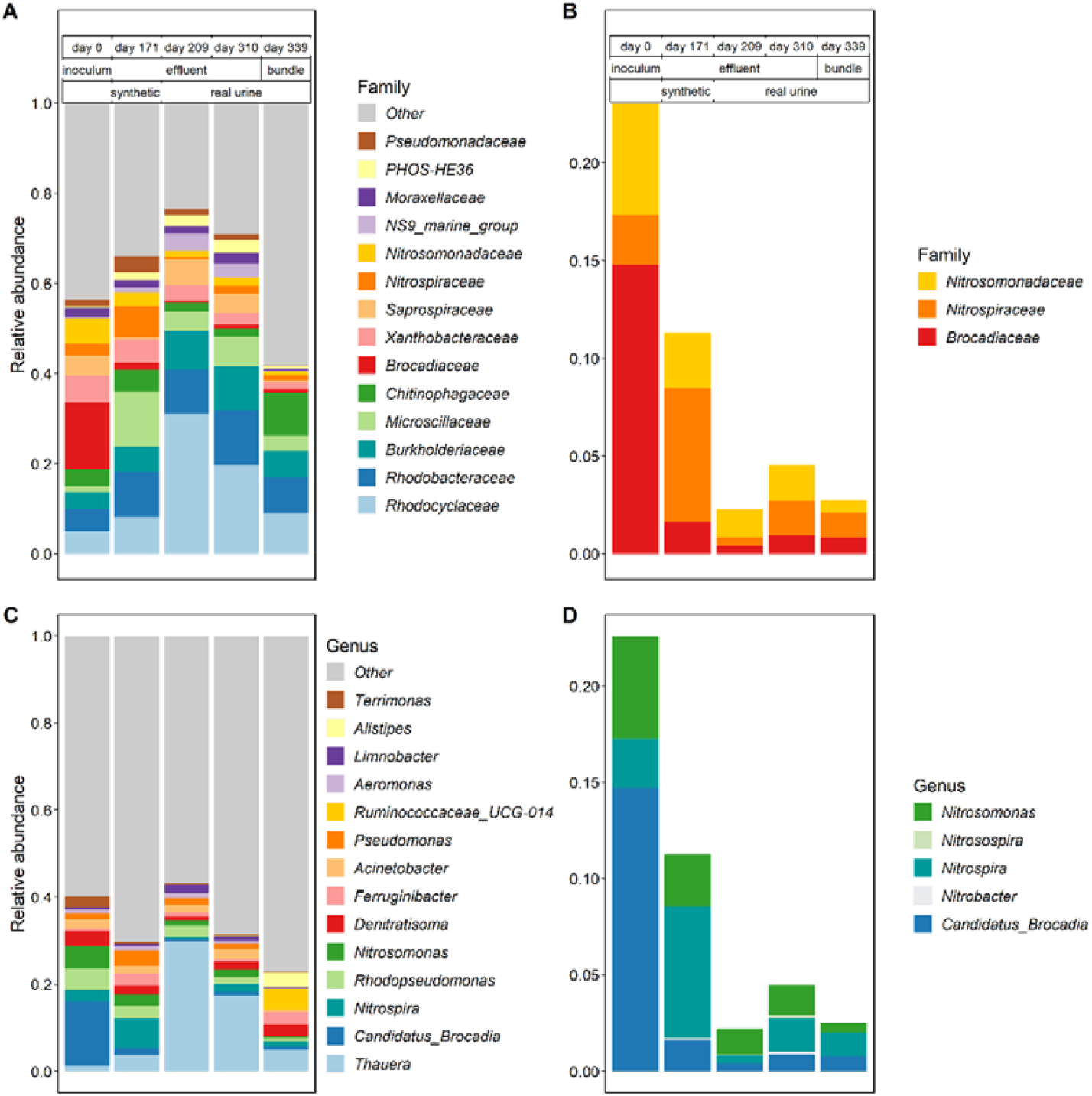
Microbial community on family level and genus level. Relative abundance is given of A) most detected families(n=15), B) families known to be involved in nitrogen removal, C) Most detected genera (n=15), D) genera known to be involved in N-removal. Shannon diversity index respectively: 3.76 (day 0), 3.67 (day 171), 3.07 (day 209), 3.35 (day 310), 4.04 (day 339).

The overall most abundant identified families were *Rhodocyclaceae, Rhodobacteraceae, Burkholderiaceae* and *Microscillaceae* (Figure 5A), mostly attributed to organoheterotroph activity. The most detected genus during operation on real urine (*Thauera*) (Figure 5C) is belong to the nitrate reducers/denitrifiers (Etchebehere and Tiedje, 2005). *Denitratisoma* has members that can execute denitratation (Ramos et al., 2016).

The community involved in nitrogen removal decreased in abundance during the operation on real urine, which contains a varied matrix of compounds to be converted. Consequently, a variety of heterotrophic species (for COD- and organic nitrogen conversion) grew into the reactor yielding a more diverse community than the inoculum. Despite the loss in relative abundance (Figure 5 B&D), N-removal was not affected throughout the operation on real urine. The diversity dropped during the operation on synthetic urine compared to the start-up phase with bioaugmentations (Shannon diversity index from 3.76 at inoculum to 3.09 direct after change to real urine), but the diversity raised during the operation on real urine to 4.04 (day 339).

### 3.5. Stratification of nitrifiers and anammox bacteria in the biofilm

The microbial stratification within the biofilms was investigated by cryosection of the biofilm and assessing the abundance of certain guilds using FISH. AerAOB occupy a 20 μm thick layer covering the surface of the hollow fibre membrane (Figure 6A) while NOB grew in unevenly spread 5-10 μm diameter clusters in the AerAOB layer. The AerAOB-NOB layer was lined by a 20 μm thick layer of non-stained bacteria (Figure 6D) that were potentially heterotrophic (facultatively anoxic) bacteria. This layer could have been preferable for microorganisms metabolizing larger organic compounds facing diffusive limitations as it is closer to the liquid bulk, despite the lower oxygen concentrations. Non-stained bacteria are also found directly at the membrane surface. Here, they might be aerobic heterotrophs that convert COD, competing with AerAOB and NOB for oxygen.

**Figure 6.**
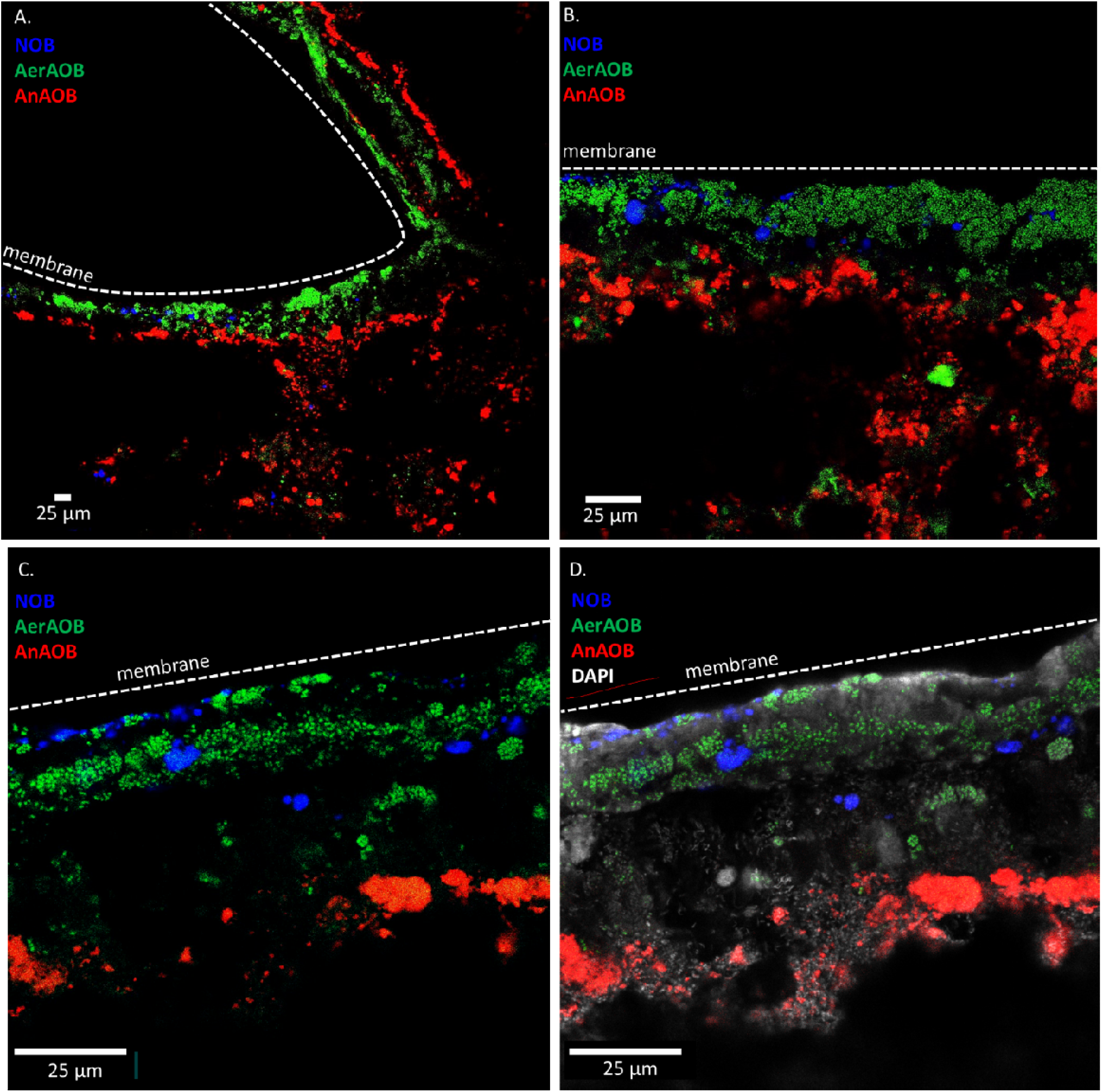
Stratification of N removal community in PN/A MABR biofilm with FISH. Blue=NOB, Green=AerAOB, Red=AnAOB, White=DAPI (all DNA), only fig D. Magnifications; A: 1:200, B:1:630, C&D:1:950.

Within the anoxic layer (average thickness of 500 μm), AnAOB were mostly observed at 40 to 100 μm from the aerated membrane. This stratification of AnAOB close to the oxic zone is most likely driven by the need for anoxic conditions and by NO_2_ limitation: NO_2_^-^ rarely accumulated, so proximity to the NO_2_^-^ producing AerAOB yielded more substrate availability compared to colonies deeper in the anoxic zone of the biofilm.

## 4. Discussion

### 4.1. Stable urine treatment with MABR-PN/A

Stable PN/A was achieved in MABR on diluted synthetic and real urine throughout an operation span of 500 days. The highest NRR of 1.6 g N L^-1^ d^-1^ (2.5 g N m^-2^ d^-1^ ) was obtained when the system was overloaded, which results in limited NRE of 60%. A trade-off between NRR and NRE was outlined, as a NRE of 90% was obtained at a lower NRR of 1.0 g N L^-1^ d^-1^ (1.5 g N m^-2^ d^-1^ ). The obtained NRRs are within the range of typical volumetric (between 0.3-2.0 mg N L^-1^ d^-1^ (Pellicer-Nàcher et al., 2010; Sliekers et al., 2003; Vlaeminck et al., 2009)) and membrane area specific (1500-2500 mg N m^-2^ d^-1^ (Gilmore et al., 2013)) rates for PN/A in sidestream conditions, at high (>25°C) temperatures. Compared to previous work of PN/A on urine, the rates and efficiencies obtained are more than double and maintained for much longer than previous work (Bürgmann et al., 2011; Zhan et al., 2022).

The superior performance of PN/A-MABR could be attributed to the nature of biofilm growth, which differs fundamentally from suspended or floccular reactors. The two mechanisms that potentially contribute to the enhanced performance of MABR are the overcapacity in anammox activity that can be maintained and the shielding effect of the biofilm. Besides during the operation at maximum NRR (1.6 g N L^-1^ d^-1^), rates were not limited by anoxic activity but by aeration/prevention of NOB activation. Therefore, any disruptions (e.g., shocks in load, COD/N fluctuations) causing partial inhibition can be easier accommodated for and will not lead to failure. The growth in biofilm retains any AnAOB that upon partial inhibition might have washed out in floccular systems (Bürgmann et al., 2011; Schielke-Jenni, 2015).

Moreover, the biofilm also shields AnAOB from exposure towards any shocks. Specifically, within the anoxic biofilm layer in MABR, the highest AnAOB presence was found to be close to the oxic layer and the membrane, further away from the liquid bulk. The thick (>500 μm) anoxic layer that was maintained in this reactor effectively shields the AnAOB-rich zone. By operating in side-stream conditions such as urine, diffusion limitations are reached later than in mainstream conditions, effectively allowing for a thicker biofilm: Given the autotrophic nature of PN/A, only three wasting events had to be performed during the operation span whilst not impacting performance.

COD-removal was considered beneficial for the working of the reactor. Firstly, heterotrophic organisms scavenge remaining oxygen in the reactor. This led to a steeper gradient of oxygen, allowing AnAOB to grow at 40 μm from the membrane surface compared to autotrophic biofilms (Gilmore et al., 2013), where the main AnAOB presence was found 400 μm from the surface under similar conditions. Secondly, heterotrophs are crucial in biofilm formation. Stabilization of urine causes precipitation and removal of divalent cations in the urine matrix (Paepe et al., 2021; Randall et al., 2016) which complicates bacterial attachment and biofilm formation (Sobeck and Higgins, 2002). *Rhodocyclaceae* was the most abundant heterotrophic family in the MABR and its members are known to be involved in the production of biopolymers such as EPS (Weissbrodt et al., 2014) and biofilm formation.

### 4.2. The crucial role of oxygen supply to suppress NOB activity and maximize N removal

Although the MABR was able to provide stable nitrogen removal on diluted urine, a combination of strategies had to be supplied to continuously supress NOB. Firstly, oxygen supply was optimized and a tipping point between TAN or NO_3_^-^ residual was reached between 30-35 min h^-1^ . Optimum NRE was reached at aeration time of 27 min h^-1^ and DO setpoint 0.05 mg O_2_ L^-1^ . Although combined control yielded almost complete NOB suppression for a short-term period of 20 days (after inoculation of a new bundle), growth over time and the absence of regular scouring created a thick biofilm (>500 μm). This resulted in limited response of the DO-setpoint control in bulk and led to eventual NOB activation. Whereas scouring would control biofilm thickness and might yield more precise oxygen supply control, it might come at the cost of AnAOB-capacity which is essential for resilience during operation on urine.

Moreover, the here described optimum aerated time oxygen is highly specific: fluctuations in influent (COD/N) and load have been reported to heavily affect PN/A performance in MABR (Bunse et al., 2020; Larsen et al., 2021). Oxygen uptake rate (OUR)-based control in combination with strictly anoxic time (OUR+timer) could be a more viable control strategy as it provides a stable measurement for oxygen demand in the reactor (thus oxygen gradient in the biofilm) that can indicate NOB activation and could work independent of biofilm thickness. Additionally, OUR-control is easy to implement given the MABR-off gas is a piped stream and its limited maintenance.

### 4.3. MABR-PN/A treatment of undiluted urine at the same the N removal efficiency

To further reduce dilution requirement for treatment, MABR-PN/A was also operated on less and undiluted urine (33%, 66%, 100%). The optimum removal efficiency of 93% was reached on 66% urine as inhibitory effects started to lower the efficiency to 85% at undiluted urine. This shows that the PN/A process in MABR could adapt at least up to 66% of urine without major performance impact. Further work should confirm the constancy of these short term observations and investigate whether any adaptation of AerAOB and AnAOB might occur that could alleviate inhibition over time as described for saline matrices in (Windey et al., 2005). In this reactor, only Candidatus Brocadia was detected as AnAOB. Possibly, limitations could be overcome at the 100% urine matrix as well by introducing halotolerant AnAOB species, such as Candidatus Kuenenia (Kartal et al., 2006).

### 4.4. Implementation perspective and effluent applications

The PN/A-MABR fits the properties of source-separated decentralized urine treatment (Garrido-Baserba et al., 2018; Rabaey et al., 2020) due to the superior energy efficiency of MABR technology (Syron et al., 2015) compared to the state of the art and the lower oxygen requirement of the PN/A compared to N/DN. Leveraging both advantages, the PN/A-MABR described here could cut energy demand from 45 MJ kg^-1^ N to ∼6 MJ kg^-1^ N, or approximately 85% energy reduction compared to conventional nitrification systems. Moreover, the optimum NRE (93%) was achieved at urine concentrations of 4 g N L^-1^ . This is a concentration typical in urine diversion systems (Maurer et al., 2003) and underlines its applicability in such systems.

About 98% of the TAN in the urine was removed during optimal reactor performance, but the effluent does still not comply with TN standards for discharge (10-20 mg N L^-1^) and reuse (10-50 mg N, mostly as N-NO_3_^-^, depending on application) (Alcalde and Gawlik, 2014; European Commission, 2014; Lavrnić et al., 2017). Therefore, the effluent can either be discharged in the sewer or it can be further upgraded with an additional treatment step e.g., treatment mixed with normally N-limited grey water (Jefferson et al., 2001) in MBR (Garrido-Baserba et al., 2018), to simultaneously remove suspended material. This could render the water re-usable for non-potable purposes.

### 4.5. Nitrogen gas production with a PN/A-MABR for regenerative life-support systems in Space

The PN/A-MABR reactor could also be implemented in regenerative biological life support systems for long-duration human spaceflights where it can produce water and N_2_ from urine. State-of-the-art urine treatment in space now consists of chemical stabilization with CrO_3_, which is one of the most toxic substances present at the international space station, followed by vacuum compression distillation (VCD) (Holder and Hutchens, 2003). The VCD has a nominal distillation efficiency of 85%, which is less than the 98% water recovery which NASA deemed necessary for long-term human space flight (Hurlbert et al., 2012). Biological urine treatment has no theoretical constraints on maximal water recovery and allows for tailored desalination step in absence of ammonium.

An added benefit of biological urine treatment is that the produced N_2_ can serve as ullage gas to manage cabin pressure. Gas losses occur due to structural leaks and extravehicular activities. The cost of these losses adds up when long-term missions are considered. A Mars-transit mission lasting 900 days can lose up to 22 – 52 kg N_2_-N depending on the ECLSS configuration (Goodliff et al., 2017). State-of-the-art methods to counteract these losses are hauling pressurized containers which pose both mass and safety constraints on the mission architecture. Using a 40 L reactor, urine of 4 astronauts could be converted to supply 65 – 180 % of the N_2_-N requirements providing additional benefits beyond water recovery.

Microgravity-compatible MABR technology has already been investigated for urine nitrification in combination with safer electrochemical stabilization (Paepe et al., 2020) and even N/DN (Landes et al., 2021), denitrification is severely limited because of the low COD/N in urine. As such, a PNA-MABR approach would be an ideal candidate to replace the VCD technology as it maximizes circularity (in terms of water and N_2_), it minimizes consumable and energy cost, and significantly leverages the autotrophic nature of PN/A as sludge production and its associated disposal can be minimized.

## 5. Conclusions

- Aeration- and carbon-efficient N removal on real urine was established in a PN/A-MABR, with a trade-off between maximum N removal rate at lower efficiency vs. lower rate at maximum efficiency.
- The MABR was able to overcome process instabilities and failure in previous PN/A treatment of urine, potentially due to the biofilm stratification that allows (1) overcapacity in AnAOB, and (2) biomass retention preventing washout and (3) shielding to COD/N ratio variations.
- Although a nitrogen removal efficiency of over 90% was reached at high rates, the delicate balance between AOB-limitation by DO and NOB-activation from high DO makes it impossible to sustain such NREs in the long term, even with advanced control strategies such as intermittent and DO-controlled aeration.
- PN/A could even be operated on less- and undiluted urine with an optimum nitrogen removal efficiency of 93% for urine dilutions at 66% (4 g N L^-1^), whilst partial inhibition was observed at pure urine (5.5 g N L^-1^), only reaching 85% removal efficiency.
- An increasingly diverse microbial community was found especially on real urine, with a small functional N-converting community dominated by *Candidatus* Brocadia, *Nitrosomonas* and *Nitrospira* members.
- PN/A in MABR can serve as energy and carbon lean alternative for urine treatment on earth in decentralized systems to remove the majority of N and C and in regenerative life support systems in space such as MELiSSA for purification of water and production of nitrogen as cabin ullage gas.

## 6. Acknowledgements

M.T. and J.D.P. were supported by a grant from the European Space Agency (ESA, contract no. 4000130179/20/NL/KML), and is part of MELiSSA project (www.MELiSSAfoundation.com). K.D.P. was funded by Ghent University (BOF/GOA 01G03122). I.S. was supported by the Research Foundation – Flanders (Fonds Wetenschappelijk Onderzoek (FWO)) postdoctoral grant 1277222N. Fluorescence microscopy work was supported by the European Research Council grant Lacto-Be 26850 of S.L. R.G. was supported by the Special Research Fund of Ghent University (grant number BOF19/STA/044) and the FWO (G052520N). The authors would like to thank Philipp Markus for his input, Dries De Laet, Thomas Chretién, and Isabel Morowa for their support with the reactor operation and Nele Van de Vliet for assistance with FISH staining.

## 7. Declaration of competing interests

The authors declare that they have no known competing financial interests or personal relationships that could have appeared to influence the work reported in this paper.

## Notes

### Competing Interest Statement

The authors have declared no competing interest.

