## Supplementary information for "Resource-efficient nitrogen removal from source separated urine with partial nitritation/anammox in a membrane aerated biofilm reactor"

### A. Process flow diagram and picture of the membrane aerated biofilm reactor

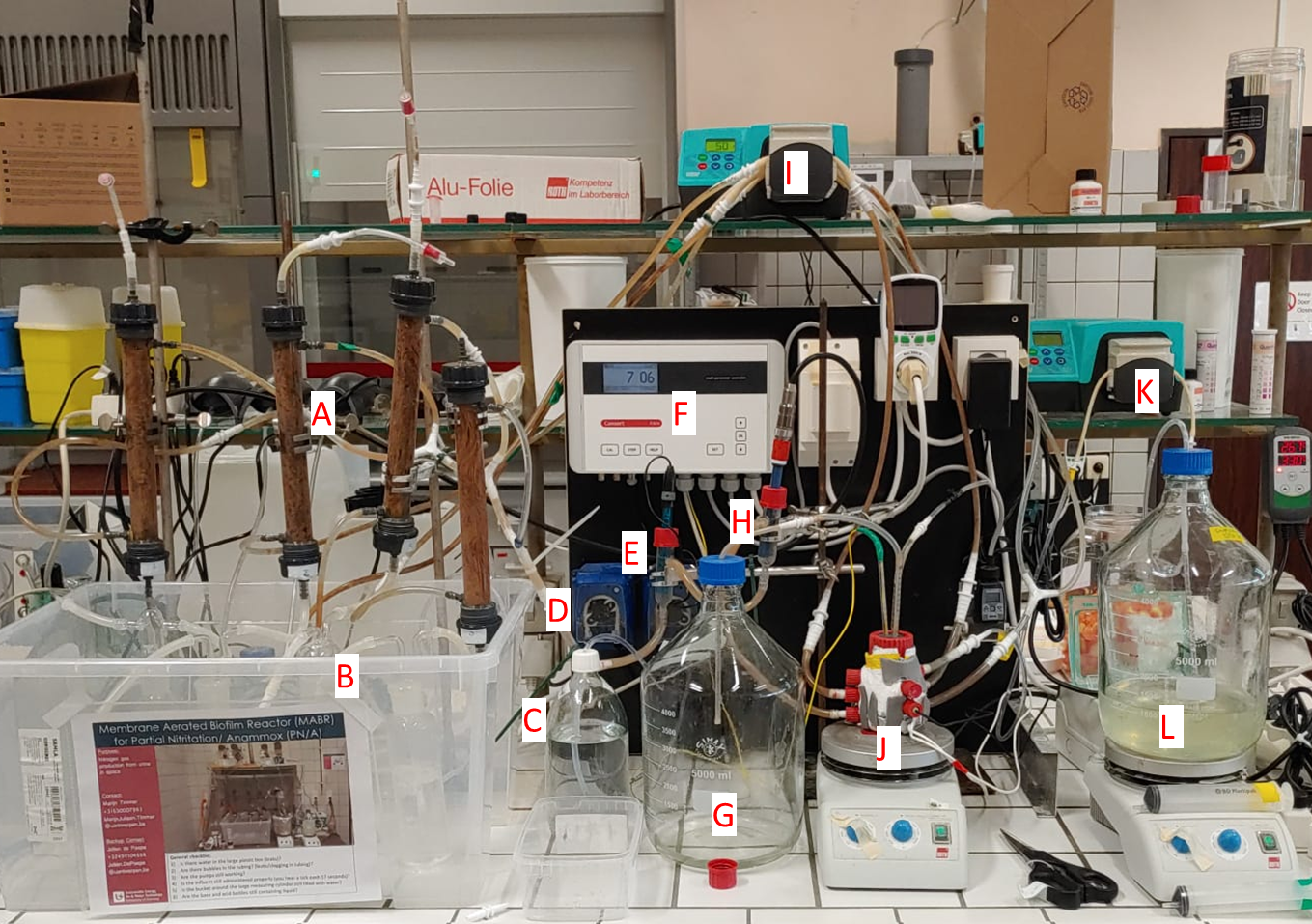

Figure S1. MABR setup in lab

A: Membrane modules	B: Washing bottles	C: HCl-storage

D: HCl-dosage pump	E: pH-flowcel	F: Controller

G: Effluent vessel	H: DO-flowcel	I: Recirculation pump

J: Mixing vessel	K: Influent pump	L: Influent vessel

### B. Feeding and aeration profiles

Table S1. Regimes used for the optimization of feeding and aeration during the whole reactor operation span

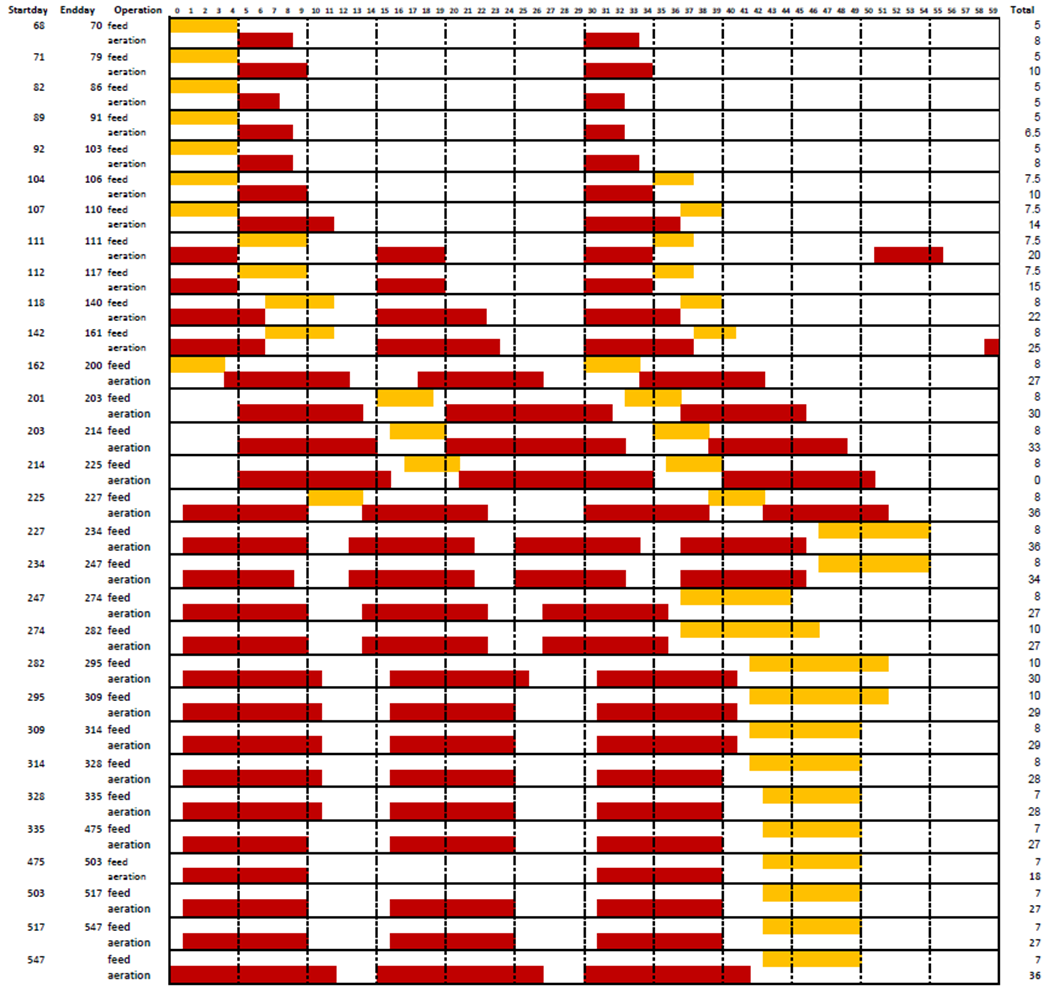

### C. Recipes of synthetic urine

Table S2. Recipes of synthetic urine

|  | **synthetic hydrolysed urine** | **synthetic urine with Na-Ac as COD** | **synthetic urine with complex COD** |
| --- | --- | --- | --- |
| **Phases used** | S-HUR | S-UR I | S-UR II-VI |
| **Compound** | concentration[g l^-1^] | concentration[g l^-1^] | concentration [g l^-1^] |
| Urea |  | 1.078 | 1.078 |
| (NH_4_)_2_SO_4_ | 3.162 | 0.790 | 0.790 |
| Creatinine |  |  | 0.462 |
| Citric acid |  |  | 0.159 |
| Hippuric acid |  |  | 0.168 |
| Na-Ac | 0.981 | 0.981 |  |
| NaHCO_3_ | 4.020 |  |  |
| KCl | 0.500 | 0.500 | 0.500 |
| KH_2_PO_4_ | 0.236 | 0.236 | 0.236 |
| CaCl_2_.7H_2_O | 0.070 | 0.070 | 0.070 |
| MgSO_4_.7H_2_O | 0.100 | 0.100 | 0.100 |

### D. Pipeline used for Miseq data

Clipping of adapter and primer sequences was done using cutadapt (v4.1). Reads with ambiguous base-calls (Ns) were first removed to accurately map short adapter and primer sequences. Paired-end reads without an exact match to the forward and reverse adapter and primer sequence were removed. Further processing of the sequencing data was done with the DADA2 R package (v1.24.0) in R (v 4.2.2) according to (Callahan et al. 2016). As first quality control step, truncation was done with a quality score cut-off (truncQ=2). Additional filtering to eliminate reads containing any ambiguous base calls or reads with high expected errors (maxEE=2,2) was performed. Reads mapping to the Phix genome were discarded as well. After dereplication, further denoising using the Divisive Amplicon Denoising Algorithm (DADA) error estimation algorithm and the sample inference algorithm selfConsist (with nbases=1e8) was done. The obtained error rates were checked and monotonicity was enforced in the model fit to deal with binned quality scores. The loess function was reweighted (log10 transformation of total transition probabilities) whilst the span was adjusted to 2. Denoised reads inferred were subsequently merged. Finally, the amplicon sequence variant (ASV) table obtained after the removal of chimera was used for taxonomy assignment. This was done with the Naive Bayesian Classifier (with an 80% minimal bootstrap confidence threshold) and the DADA2 formatted Silva v132 and RDP trainset 16 reference taxonomy according to (Quast et al. 2013; Wang et al. 2007).

### E. Fish probes used

Table S3. FISH probes used for identification of Aer-AOB, AnAOB, and NOB

| **group** | **probe** | **target** | **sequence** | **reference** |
| --- | --- | --- | --- | --- |
| Aer-AOB | Nso1225 | Betaproteobacterial AerAOB | CGCCATTGTATTACGTGTGA | Mobarry et al. 1996 |
|  | Nso190 | Betaproteobacterial AerAOB | CGATCCCCTGCTTTTCTCC | Mobarry et al. 1996 |
| AnAOB | Amx368 | Anammox bacteria | CCTTTCGGGCATTGCGAA | Schmidt et al. 2003 |
|  | Amx820 | *“Candidatus Brocadia”* and *“Candidatus Kuenenia”* | AAAACCCCTCTACTTAGTGCCC | Schmidt et al. 2003 |
| NOB | NIT3 + competitor | Nitrobacter spp. | CCTGTGCTCCATGCTCCG | Wagner et al. 1996 |
|  | Ntspa662 + competitor | Nitrospira spp. | GGAATTCCGCGCTCCTCT | Daims et al. 2001 |

### F. Area specific nitrogen removal rate

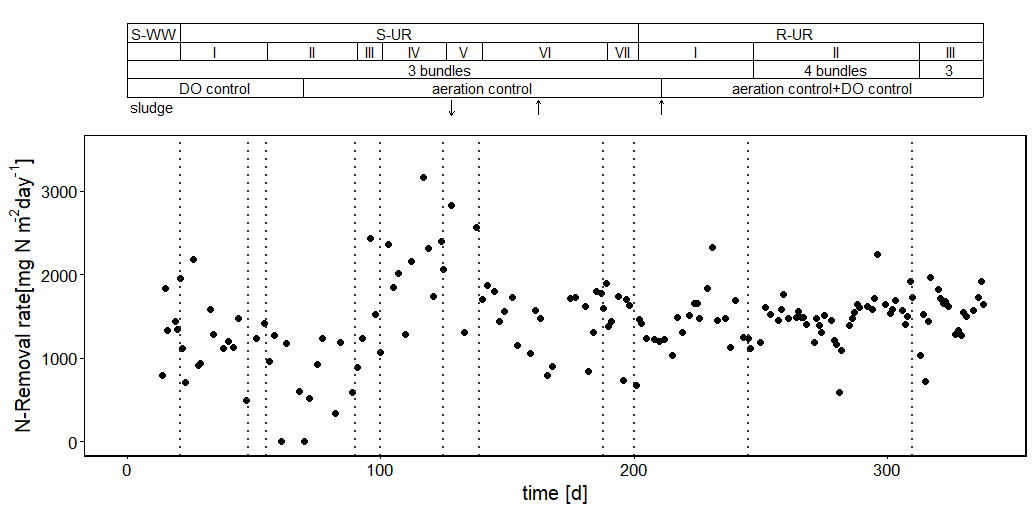

Figure S3. Area specific nitrogen removal rate

### G. Nitrogen conversion activity in the biofilm and bulk

Table S4. Maximum anoxic activity in the different HF modules based on anoxic activity tests. Activity tests were performed with a duration of 4 h. Spikes of 100 mg N L^-1^ NH_4_^+^ and N-NO_2_^-^ were supplied

| **Reactor part** | **days of operation**  **[d]** | **NO_2_^-^ removal**  **[mg N L^-1^ h^-1^]** | **Total anoxic-activity**  **[mg N L^-1^ h^-1^]** | **Ratio NO_2_^-^/NH_4_^+^** |
| --- | --- | --- | --- | --- |
| Aeration module 1 | 270 | 33.0 | 60 | 1.22 |
| Aeration module 2 | 270 | 30.1 | 53 | 1.30 |
| Aeration module 3 | 270 | 29.6 | 52 | 1.32 |
| Aeration module 4 | 30 | 17.0 | 18 | 17 |
| Recirculation hardware | 270 | 4.3-8.6 | 12-18 | 0.56-0.89 |

### H. Performance per operational phase in MABR

| **Phase** | |  | Day | N loading | NRR | NRE | TkNRE |
| --- | --- | --- | --- | --- | --- | --- | --- |
|  |  | |  | *[g L^-1^ d^-1^]* | *[g L^-1^ d^-1^]* | *[-]* | *[-]* |
| **SH-UR** |  | | 1-4 | 0.12-0.96 | 0.0-0.82 | 0.80±0.15 | 0.98+0.01 |
| **S-UR** | **I** | | 21-55 | 1.08-1.44 | 0.72±0.12 | 0.56±0.09 | 0.98+0.02 |
|  | **II** | | 55-90 | 0.74-1.26 | 0.74±0.12 | 0.74±0.12 | 0.90+0.09 |
|  | **III** | | 90-100 | 1.54 | 1.25±0.35 | 0.81±0.21 | 0.93+0.06 |
|  | **IV** | | 100-125 | 1.79 | 1.39±0.38 | 0.78±0.22 | 0.85+0.14 |
|  | **V** | | 125-139 | 2.53 | 1.94±0.34 | 0.72±0.13 | 0.72+0.15 |
|  | **VI** | | 139-188 | 1.16-1.54 | 1.06±0.22 | 0.68±0.18 | 0.70+0.15 |
|  | **VII** | | 188-200 | 1.16-1.54 | 1.00±0.15 | 0.66±0.12 | 0.72+0.15 |
| **R-UR** | **I** | | 200-245 | 1.16-1.54 | 0.93±0.13 | 0.71±0.10 | 0.85+0.14 |
|  | **II** | | 245-310 | 1.1 | 1.01±0.14 | 0.91±0.12 | 0.93+0.05 |
|  | **III** | | 310-335 | 1.26-1.37 | 0.97±0.24 | 0.75±0.03 | 0.88+0.09 |
|  | **IV** | | 554-560 | 0.6-0.82 | 0.65±0.08 | 0.86±0.01 | 0.94±0.01 |
|  | **V** | | 566-570 | 0.6-0.82 | 0.59±0.04 | 0.89±0.04 | 0.99±0.01 |
|  | **VI** | | 570-602 | 0.6-0.82 | 0.73±0.06 | 0.90±0.07 | 0.98±0.01 |

Table S5. Performance per operational phase in MABR. Summarized Nitrogen removal rate, efficiency and kjeldahl nitrogen removal efficiency

### I. Daily performance PN/A in MABR on undiluted urine

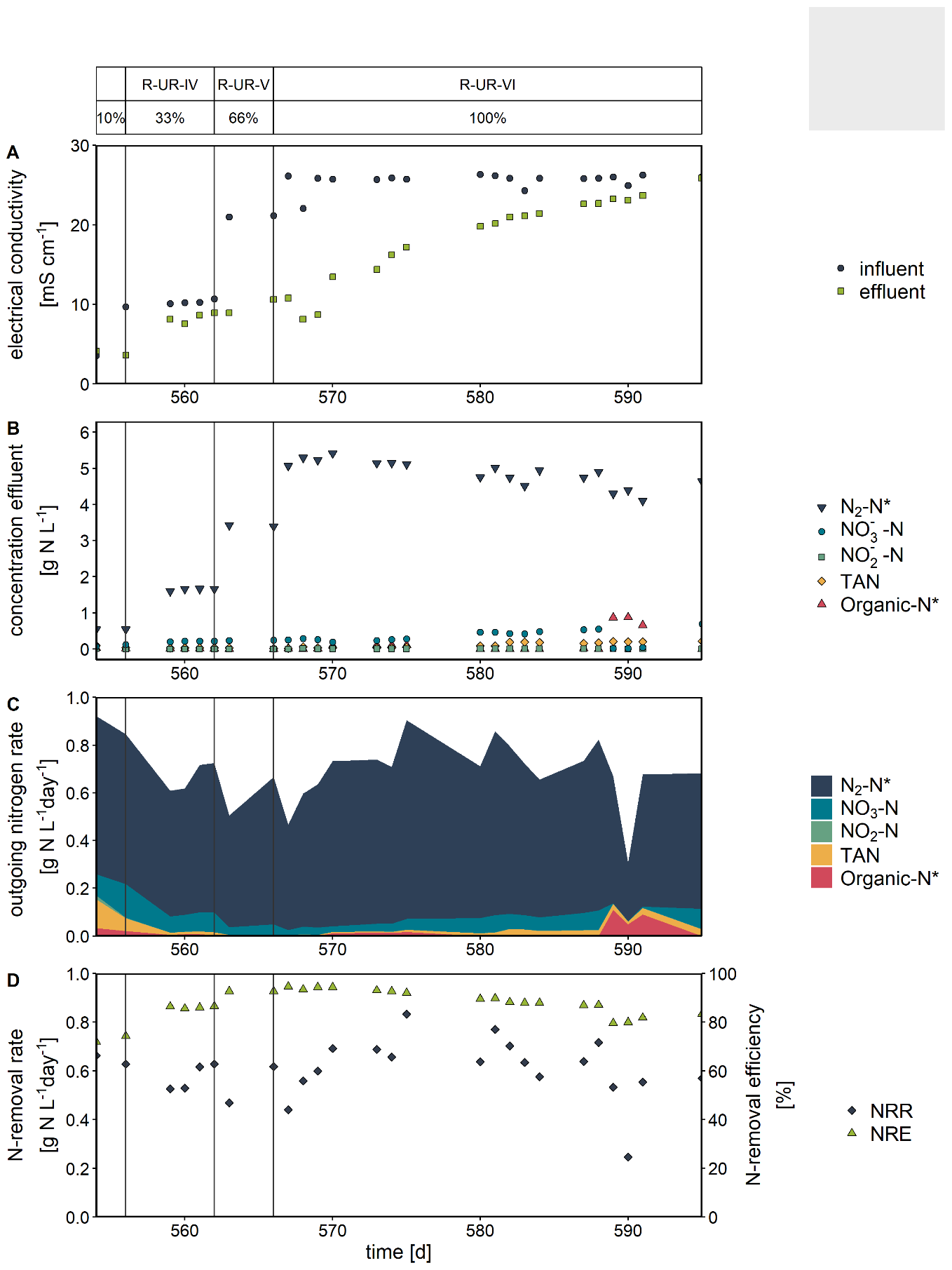

Figure S4. Performance of MABR on less- and undiluted urine. A) Conductivity in- and effluent, B) N concentrations effluent, C) Outgoing nitrogen rate and D) Nitrogen removal rate& efficiency on 33%, 66%, and 100% of urine fraction. * Derived from indirect measurements.

### J. AnAOB-activity at different salinities

N - removal rate [g N2-N L-1 d-1]

| Electrical Conductivity [mS cm-1] | N-removal per day per liter | N-removal per day per liter |
| --- | --- | --- |
| 4.17 | 1227.75 | 1227.75 |
| 9.88 | 1264.08 | 1264.08 |
| 13.01 | 1063.38 | 1063.38 |
| 19.04 | 907.845 | 907.845 |

Figure S5. AnAOB activity at increasing salinity corresponding to matrices at 33%, 66% and 100% urine matrix. Data presented is the result from anoxic activity tests done in duplicate.
